# Inhibition of Aromatase by Hops, Licorice Species, and their bioactive compounds in Postmenopausal Breast Tissue

**DOI:** 10.1101/2022.05.06.490985

**Authors:** Atieh Hajirahimkhan, Caitlin Howell, Shao-Nong Chen, Susan E. Clare, Guido F. Pauli, Seema A. Khan, Judy L. Bolton, Birgit M. Dietz

**Affiliations:** Division of Breast Surgery, Robert H. Lurie Comprehensive Cancer Center, Feinberg School of Medicine, Northwestern University, Chicago, IL; Department of Physiology and Biophysics, College of Medicine, University of Illinois at Chicago, Chicago, IL; UIC Center for Botanical Dietary Supplements Research, Pharmacognosy Institute and Department of Pharmaceutical Sciences, College of Pharmacy, University of Illinois at Chicago, Chicago, IL

**Keywords:** Aromatase, botanicals, breast cancer prevention, hops, licorice

## Abstract

Breast cancer risk continues to rise post menopause. Endocrine therapies are employed to prevent postmenopausal breast cancer in high-risk women. However, their adverse effects have reduced acceptability and overall success in cancer preventionatural products such as hops (*Humulus lupulus*) and pharmacopoeial licorice (*Glycyrrhiza*) species, which are used for managing menopausal symptoms, have demonstrated estrogenic and chemopreventive properties. Their beneficial effects on aromatase activity and expression as important factors in postmenopausal breast carcinogenesis are understudied. The presented data show that *Gycyrrhiza inflata* (GI) has the highest aromatase inhibition potency among these plants. Moreover, phytoestrogens such as 8-prenylnaringenin from hops as well as liquiritigenin and 8-prenylapigenin from licorice are shown to be potent bioactives, in line with computational docking studies. 8-Prenylnaringenin, GI extract, liquiritigenin, and licochalcone A all suppress aromatase expression in postmenopausal women’s breast tissue. Collectively, these data suggest that these natural products may have breast cancer prevention potential for high-risk postmenopausal women.

## Introduction

Breast cancer is the most common cancer and the second leading cause of cancer mortality in women, worldwide (1). Studies have frequently shown that the majority of breast cancers are estrogen dependent or estrogen receptor positive (ER+) malignancies (2). ER+ breast cancers are also the most common type in postmenopausal patients, despite the absence of circulating ovarian estrogens (2). It is posited that estrogen biosynthesis in the adipose tissue, especially in the breast adipocytes plays an important role in sustaining exposure to estrogens and the development of ER+ breast tumors after menopause (3). Estrogen biosynthesis, whether in the ovaries or other organs/tissues, involves the conversion of the androgens, testosterone and androstenedione, to their respective estrogens, 17β-estradiol (E_2_) and estrone (E_1_), by the catalytic action of aromatase (CYP19A1), a member of the CYP450 enzymes family (Figure 1). While additional steps catalyzed by enzymes such as aldoketoreductase 1C3 (AKR1C3), which converts E_1_ to E_2_, are important in estrogen biosynthesis, aromatase is the rate determining enzyme. Thus, itsregulation and activation plays a key role in estrogen production and carcinogenesis (4).

**Figure 1:**
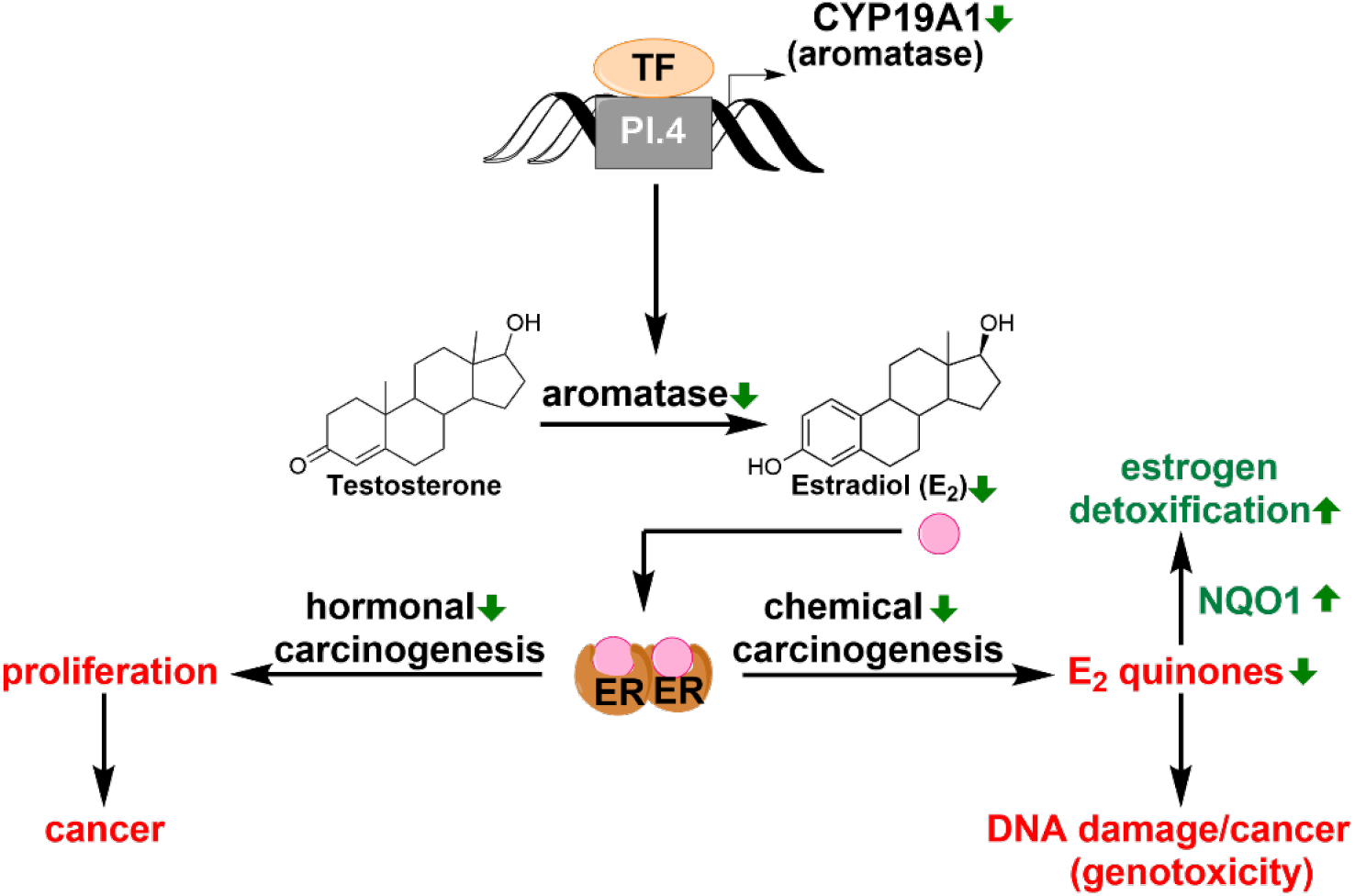
Aromatase inhibition/suppression and breast cancer prevention Aromatase is the rate limiting enzyme in estrogen biosynthesis. Suppression of its expression as well as inhibition of its activity decrease estrogen production, leading to reduced proliferation and tumor prevention, lowered hormone dependent proliferation, and reduced formation of genotoxic estrogen metabolites.

Endocrine therapies such as aromatase inhibitors (AIs, Figure 2A) have been considered as treatment options for postmenopausal women with ER+ breast cancer and premenopausal ER+ breast cancers in women with oophorectomy and ovarian suppression (5). AIs such as letrozole (non-steroidal) and exemestane (steroidal) can effectively suppress estrogen levels in breast cancer patients and enhance survival (6). In addition, these medications have been used as risk reducing agents in postmenopausal women at high risk for developing breast cancer (2). While these drugs can reduce the risk by 50 – 65 %, they have had minimal impact on lowering the breast cancer incidence in the population; mainly because less than 1% of women who would potentially benefit from such interventions report taking them (2,5,7). This is a result of their limited compliance due to side effects and to postmenopausal women’s reluctance to accept drugs for prevention purpose (2,5,7). Alternative interventions are needed with adequate efficacy yet lower toxicity that would lead to greater acceptability and better incorporation in the women’s lives.

**Figure 2:**
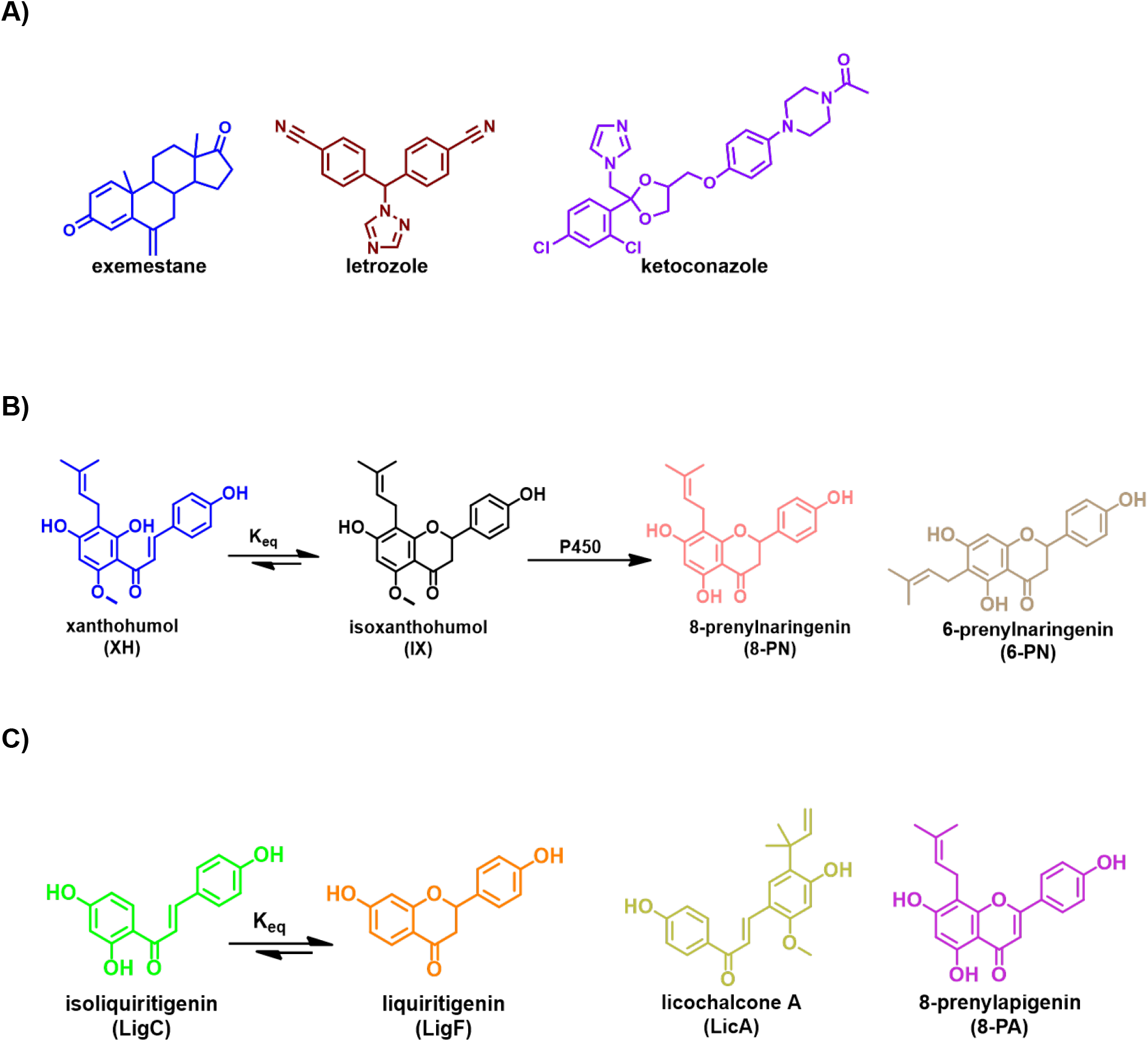
Chemical structures of A) aromatase inhibitors, B) hops bioactive compounds, and C) licorice bioactive compounds.

In search of such option, we focused on botanicals such as hops (*Humulus lupulus*) and pharmacopoeial licorice species (*Glycyrrhiza species*), all of which are widely used by women for managing menopausal symptoms (8,9). Being used as dietary supplements, these interventions are often part of daily intake and are used for extended periods of time (8,9). Independent of their efficacy in relieving menopausal discomfort, hops and licorice products have been reported to exhibit various potentially chemopreventive effects such as the suppression of estrogen genotoxic metabolism, enhancement of benign estrogen metabolism, induction of detoxification enzymes, prevention of DNA damage, as well as suppression of inflammation (8,10–24). Considering the enhanced acceptability among postmenopausal women who are at greater risk for developing breast cancer, understanding the breast cancer prevention potential of these agents could lead to more successful prevention strategies (8).

Therefore, the current study systematically evaluated the effects of hops, and the three *Glycyrrhiza* species (*G. glabra*, GG; *G. uralensis*, GU; *G. inflata*, GI) that are still used interchangeably as “licorice” botanical dietary supplements, and their major bioactive constituents in inhibiting aromatase. The study also compared the binding of bioactive marker compounds of hops and licorice species to aromatase using computational approaches. Additionally, changes in aromatase expression in postmenopausal women’s breast tissue microstructures exposed to these botanicals and their bioactive compounds were studied.

The outcomes showed that hops and licorice species as well as their bioactive compounds have the potential to limit estrogen biosynthesis through the direct inhibition of aromatase and via suppression of its expression in the breast tissue of high-risk women. Future studies are warranted to further establish the *in vivo* aromatase inhibitory effects of these natural products and their potential clinical utility in breast cancer prevention for high-risk postmenopausal women.

## Material and Methods

### Chemicals and Materials

All chemicals and reagents were purchased from Sigma-Aldrich (St. Louis, MO), unless otherwise indicated. MammoCult media kit, heparin, and hydrocortisone were purchased from Stem Cell Technologies (Vancouver, BC. Canada). F12-K nutrient mix (Kaighn’s) medium was acquired from Gibco (Dublin, Ireland). Fetal bovine serum (FBS) was purchased from Atlanta Biologicals (Norcross, GA). Collagenase I was purchased from Sigma-Aldrich (St. Louis, MO). Isoliquiritigenin (LigC) and liquiritigenin (LigF) were acquired from ChromaDex (Irvine, CA). 8-prenylapigenin (8-PA) was purchased from Ryan Scientific Inc. (Mount Pleasant, SC). Licochalcone A (LicA), 8-prenylnaringenin (8-PN), and 6-prenylnaringenin (6-PN) were acquired from Sigma-Aldrich (St. Louis, MO) and their purity assessed independently. Xanthohumol (XH) was isolated from hops (*Humulus lupulus*) as described previously (25). CYP19/MFC high throughput screening kit was purchased from Corning (Corning, NY). Direct-Zol RNA prep kit was acquired from Zymoresearch (Irvine, CA). TRIzol was obtained from Invitrogen (Waltham, MA). PCR reagents, primers, and master mix were purchased from Integrated DNA Technologies (Coralville, IA).

### Preparation and characterization of plant extracts

The ethanol extract of botanically authenticated strobili of *Humulus lupulus* was dispersed on diatomaceous earth and extracted with CO_2_ yielding spent hops. The spent hop extract was obtained from Hopsteiner (Mainburg, Germany, and New York, NY) and standardized to the prenylated polyphenol marker compounds 6-PN, 8-PN, IX, and XH as previously described (26,27). Briefly, standardization involved characterization by LC-UV, LC-MS/MS, and quantitative ^1^H NMR (qHNMR). The same extract has been used in a Phase I clinical trial in postmenopausal women (28). The concentrations of the four marker compounds in this extract were 1.2% 6-PN, 0.33% 8-PN, 0.99% IX, and 32% XH. The extracts of the three different licorice species (*Glycyrrhiza glabra* L., *G. uralensis* Fisch. ex DC., and *G. inflata* Batalin, Fabaceae) were the chemically characterized methanol extracts of the respective dried and DNA-identified licorice root powders, as described previously (12,29–31). The purity of the investigated pure compounds was determined by quantitative 1D ^1^H NMR using the 100% method (32) and yielded the following purity percentages (in % w/w): LicA 96.1% (ratio trans/cis = 93/7), LigF 96.6%, LigC 98.6%,, 8-PN 95.9%, 6-PN 98.5%, 8-PA 98.8%, and XH 96.5%.

### Aromatase inhibition assay

To evaluate the aromatase inhibitory potential of hops, licorice species, and their bioactive compounds, the Corning^®^ Supersomes™ P450 Inhibition Kit CYP19/MFC high throughput inhibition assay was used. The protocol of the kit, which is based on the effects of the tested agents on the conversion of the fluorescent substrate, 7-methoxy-4-trifluoromethyl coumarin (MFC) to 7-hydroxy-4-trifluoromethyl coumarin (HFC) by aromatase, was followed. Briefly, in a 96 well plate, NADPH cofactor mix containing cofactors, G6PDH, control protein, and water were incubated with various concentrations of the test agents and ketoconazole (positive control) in 37°C for 10 min. Then the enzyme/substrate mixture containing the aromatase enzyme and MFC were added to the pre-incubated mixture and were incubated at 37°C for 30 min before stopping the enzymatic reaction with the stop solution. The fluorescent signal was evaluated at the Excitation/Emission of 409 nm/530 nm and was corrected for the potential innate fluorescence of the tested extracts and compounds.

### In silico docking analysis

In silico docking analysis was performed to investigate the interactions of the isoflavones with the aromatase-heme complex. The binding site of heme-bound human placental aromatase in complex with androstenedione was obtained from the Protein Data Bank (PDB ID: 3EQM) and uploaded to Molecular Operating Environment (MOE; Chemical Computing Group, version 2016.0208). All water molecules and androstenedione were removed, and the MOE QuickPrep function optimized the aromatase-heme complex with standard settings, which included structural error correction, partial charges calculation, hydrogen addition (Protonate3D), and a minimization of pocket and ligand residues within 8 Å of the ligand. The resulting complex was selected as receptor for all docking studies. Compounds were docked with triangle matcher placement with London dG scoring and induced fit refinement with GBVI/WSA dG scoring. The images are shown with the docked compound in its best-fit position with pi bonds shown as dashed lines. Molecular surface showing hydrophobicity and lipophilicity of the binding site was generated by MOE (supplementary data). 2D ligand interaction diagrams were generated by MOE ligand interaction function (Supplementary Data).

### Human breast microstructure preparation and treatment

Institutional Review Board (IRB) approval (IRB STU00202331) was obtained from Northwestern University prior to consenting patients and collecting samples. To prepare breast microstructures the protocol of Brisken’s group was used with some minor modifications (33,34). The unaffected contralateral breast tissues of postmenopausal women who underwent bilateral mastectomy due to incident unilateral breast cancer were obtained while fresh and diced to approximately 5 mm pieces before digesting at 37°C with 2% collagenase I in F12-K nutrient mix (Kaighn’s) medium, overnight. After the completion of digestion, the tissue was centrifuged at 250 x g for 5 min and the supernatant was discarded. The pellet was washed with phosphate buffered saline and was mixed and cultured in MammoCult media supplemented with 0.2% heparin and 0.5% hydrocortisone. After 24 h incubation at 37°C the treatments were mixed with fresh MammoCult media and added to the microstructures.

### RNA preparation and qPCR

After 24 h incubation with the treatments, the microstructure pellet was prepared and washed with HBSS. Trizol was added to each pellet and the protocol of Direct-Zol RNA prep kit was used to extract RNA. The protocol included a step of DNAse I treatment for removing DNA contamination of the RNA preparations. RNA was eluted in nuclease free water and the concentration was measured using Nano Drop 2000 spectrophotometry (Thermo Fisher). The Integrated DNA Technologies (IDT) protocol, pre-made aromatase (*CYP19A1*) gene primer, and the pre-made housekeeping *RPLP0* gene primer was used to perform qPCR using QuantStudio 12K Flex Real Time PCR system.

### Statistical Analysis

Data were analyzed using GraphPad Prism 9 and were expressed as the means ± SD followed by unpaired t-test to express the difference between the two groups. A p-value of less than 0.025 was considered statistically significant.

## Results

### Hops and licorice species as well as their major bioactive polyphenolic constituents inhibit aromatase

Hops and the three licorice extracts (GG, GU, and GI) inhibited aromatase dose dependently (Figure 3). GI was the most potent extract with an IC_50_ value of 1 ± 0.14 μg/mL followed by GG (IC_50_ = 2.8 ± 0.12 μg/mL), GU (IC_50_ = 3.4 ± 0.12 μg/mL), and hops (IC_50_ = 4.9 ±0.10 μg/mL) (Figure 3). Among the tested bioactive marker compounds from hops (Figure 4A), 8-PN (Figure 2B) was the most potent aromatase inhibitor with an IC_50_ value of 50 nM (Table 1), which was in the range of inhibitory potency of known aromatase inhibitors such as letrozole (IC_50_ ≈ 10 - 20 nM) (35). The rank order of IC_50_ values for the compounds from hops was 8-PN (50 nM) >> XH (4.3 μM) > 6-PN (7.4 μM) (Table 1, Figure 4A). Among the licorice compounds (Figure 2C, Figure 4B), LigF exhibited the highest inhibitory potency with an IC_50_ value of 430 nM, followed by 8-PA with an IC_50_ value of 590 nM. The rank order of IC_50_ values for the compounds from the three licorice species was LigF ≥ 8-PA > LicA (3.2 μM) ≥ LigC (4 μM) (Table 1). It should be noted that LicA is a marker compound specific for GI that has not been found in other species, whereas 8-PA occurs in multiple *G*. species but is more concentrated in GI compared to other species (29,36,37).

**Figure 3:**
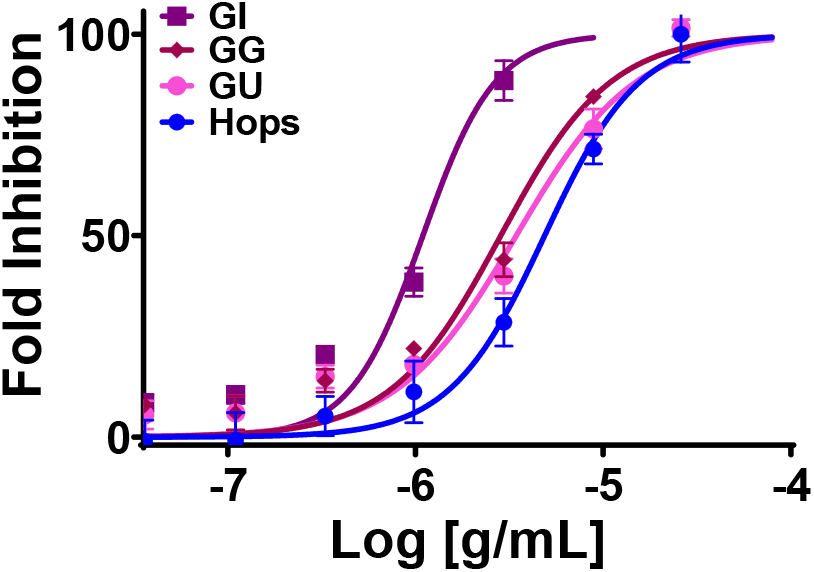
GI exhibits a more pronounced aromatase inhibition effect compared to GG, GU, and hops extracts. Aromatase supersomes were incubated with various concentrations of GI, GG, GU, and hop extracts for 40 min at 37 °C. Formation of a fluorescent metabolite was measured at Ex/Em of 409 nm/530 nm. Data represents mean ± SD of at least three independent measurements. Ketoconazole was used as the positive control.

**Figure 4:**
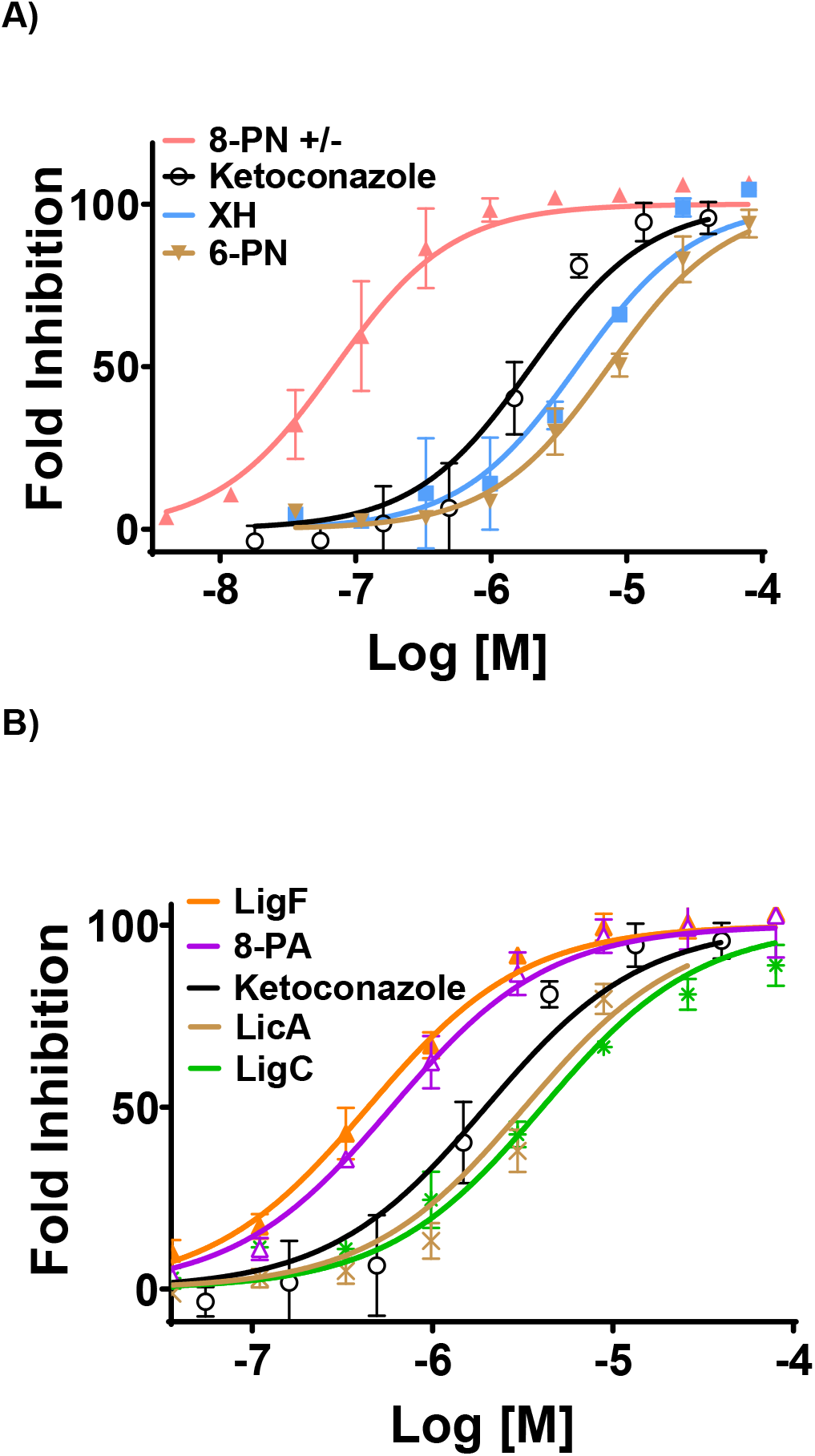
Phytoestrogens from hops and licorice exhibit the most pronounced aromatase inhibition effect compared to the other evaluated compounds. Aromatase supersomes were incubated with various concentrations of compounds from A) hops: 8-PN, XH, 6-PN and B) licorice: LigF, 8-PA, LicA, LigC for 40 min at 37 °C. Formation of a fluorescent metabolite was measured at Ex/Em of 409 nm/530 nm. Data represents mean ± SD of at least three independent measurements. Ketoconazole was used as the positive control.

**Table 1.** Aromatase inhibition and estrogen receptor subtype (ERα, ERβ) binding affinities of compounds from hops and licorice species.

| Compound | Aromatase inhibition<br>(IC<sub>50</sub>, μM) | ERα binding<br>(IC<sub>50</sub>, μM) | ERβ binding<br>(IC<sub>50</sub>, μM) |
| --- | --- | --- | --- |
| ketoconazole | 1.97 ± 0.1 | N/A | N/A |
| xanthohumol (XH) | 4.3 ± 0.6 | N/A | N/A |
| 8-prenylnaringenin (8-PN) | 0.05 ± 0.004 | 0.060 ± 0.01<sup>a</sup> | 0.16 ± 0.06<sup>a</sup> |
| 6-prenylnaringenin (6-PN) | 7.4 ± 0.3 | N/A | N/A |
| isoliquiritigenin (LigC) | 4.0 ± 0.3 | 16 ± 1<sup>b</sup> | 7.8 ± 0.1<sup>b</sup> |
| liquiritigenin (LigF) | 0.43 ± 0.02 | > 200<sup>b</sup> | 7.5 ± 0.5<sup>b</sup> |
| 8-prenylapigenin (8-PA) | 0.59 ± 0.08 | 0.1 ± 0.05<sup>a</sup> | 0.03 ± 0.01<sup>a</sup> |
| licochalcone A (LicA) | 3.2 ± 0.1 | N/A | N/A |

Data is presented as mean ± SD of at least three independent measurements. <sup>a</sup>J Agric Food Chem. 2020, 68: 10651-10663. <sup>b</sup> PLoS One, 2013, 8: e67947.

Overall, the most potent aromatase inhibitor compounds with the inhibitory potencies in the nanomolar range were the phytoestrogens, 8-PN from hops followed by LigF, and 8-PA from licorice (Table 1). On the other hand, the non-estrogenic bioactive constituents such as XH and 6-PN from hops, as well as LicA and LigC from licorice did not exhibit aromatase inhibition effects at nanomolar concentrations (Table 1).

### Phytoestrogens from hops and licorice bind to aromatase similarly to letrozole

The *in vitro* aromatase inhibition results (Figures 4A and B) along with published data (30,38) regarding the estrogenic efficacy of the compounds from hops and licorice (Table 1) suggested that the greater potency of phytoestrogens in aromatase inhibition might be due to a better binding to the aromatase binding pocket compared to the non-phytoestrogenic bioactive compounds. Therefore, the chemical structures of the three phytoestrogens, 8-PN from hops as well as LigF and 8-PA from licorice, along with the non-estrogenic compounds such as XH and 6-PN from hops as well as LicA and LigC from licorice species were docked *in silico using a* model of aromatase and compared to the binding of known aromatase inhibitors. The phytoestrogens with high binding affinity bound in a similar location to the native ligand, androstenedione (A4) (Figure S1A), letrozole (Figure 5A and Figure S1B), and exemestane (Figure S1C), which all interact with heme (39). One of letrozole’s nitrile adjacent rings is located in a hydrophobic channel of the binding pocket (Figure S1B). The hydrophobic prenyl group of the best binder, 8-PN, and the third best binder, 8-PA, extends in this same hydrophobic channel (Figure 5B and Figure 5C), contributing to their binding affinity. In its conformation with best binding affinity, 8-PA does not directly bind to heme in its most energetically favorable position (Figure 5C). However, 8PA does bind to heme in a less energetically favorable conformation (data not shown). If heme binding is important to aromatase inhibition, this less favorable conformation is a potential reason for its lower binding affinity. LicA binds within the binding pocket but does not bind to heme in any of its top ten most energetically favorable poses (Figure S2A). Although 6-PN did bind to the heme group (Figure S2B), its prenyl group did not fall within the hydrophobic pocket (Figure S3A). XH and LigC did not bind within the binding pocket (data not shown).

**Figure 5:**
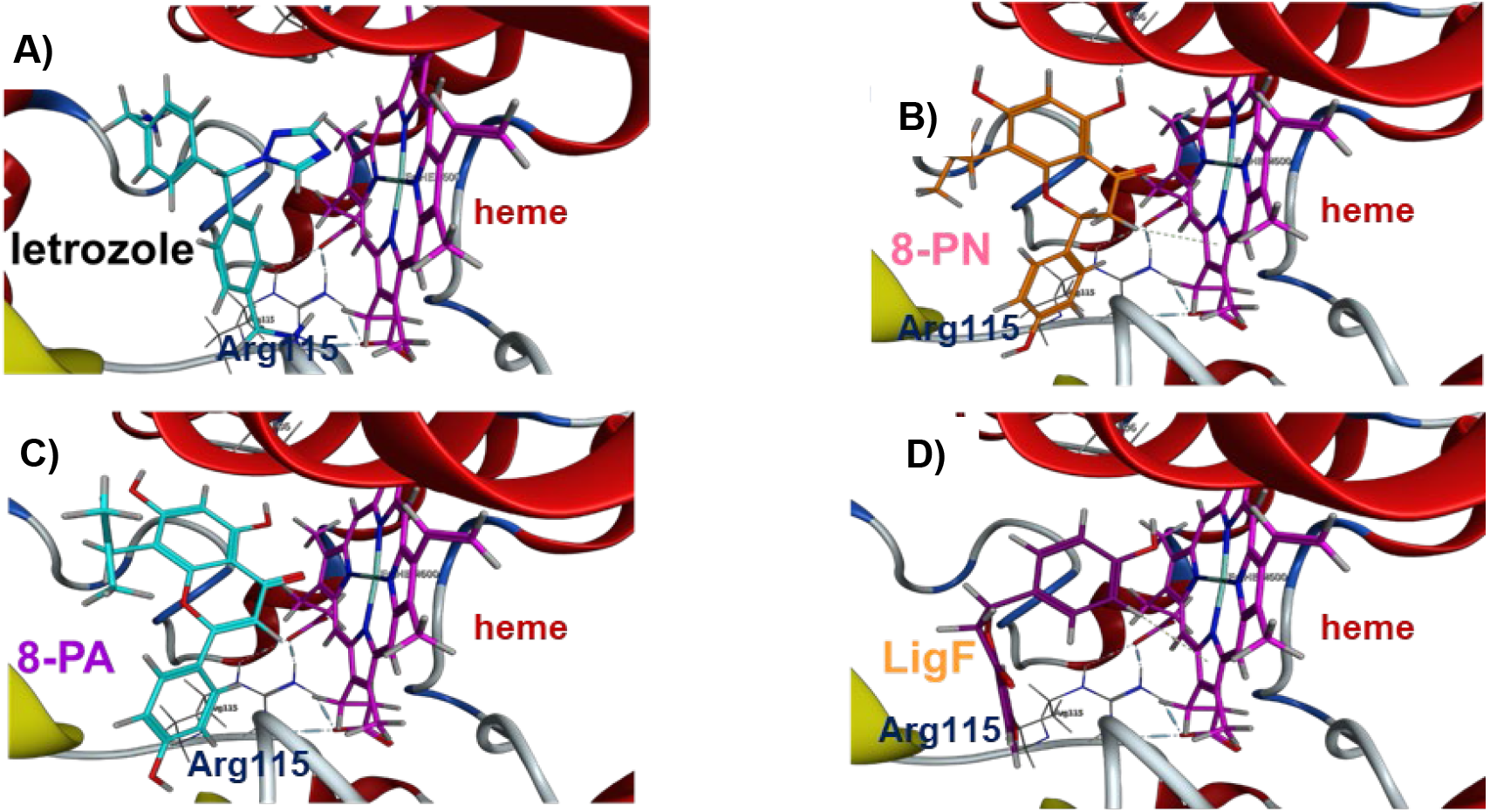
Phytoestrogens bind the active site of aromatase similar to letrozole, an established aromatase inhibitor. A) letrozole (light blue), B) 8-PN (orange), C) 8-PA (light blue), and D) LigF (dark purple) docked into aromatase (ribbon)-heme (fuchsia) complex in its most energetically favorable conformation.

### Hops, licorice, and their bioactive polyphenols suppress aromatase expression in the breast tissue of high-risk postmenopausal women

qPCR analysis of *CYP19A1* using the RNA extracted from the breast microstructures of high-risk postmenopausal women showed that the hops extract did not significantly change the base line expression levels of *CYP19A1*. However, the hop phytoestrogen, 8-PN, suppressed *CYP19A1* expression by nearly 40% in 5 subject’s tissues (p < 0.025) when compared to the vehicle treated samples. In six subject’s tissues, GI extract suppressed baseline *CYP19A1* expression by nearly 30% (p < 0.025). LigF, the key phytoestrogen from licorice, showed similar suppression levels (25%, p< 0.05), and LicA, a GI marker compound, suppressed baseline *CYP19A1* expression by nearly 40% in the six subject’s tissues (p < 0.025).

## Discussion

Breast cancer risk continues to rise after menopause despite the drastic decline in circulating ovarian estrogen (40). Menopause is associated with enhanced adiposity, which is known to lead to chronic low-grade inflammation, resulting in elevated local estrogen production in the breast stromal pre-adipocytes (3,41). Chemoprevention strategies such as AIs and selective estrogen receptor modulators (SERMs) have shown promising results in preventing breast cancer development in high-risk postmenopausal women (5,7). However, the majority of healthy but at-risk postmenopausal women refuse to take such drugs for the purpose of preventing breast cancer (42,43). Additionally, SERMs and AIs have side effects ranging from lowered quality of life to osteoporosis and cancer (42,44). These factors reduce the acceptability of the current chemoprevention interventions and as such, alternative options are needed to protect these high-risk postmenopausal women against breast cancer.

Many postmenopausal women use botanicals such as hops and licorice species (GG, GI, GU), for relieving menopausal symptoms such as hot flashes (9) and take these remedies for extended time periods as part of their daily diet, often without consulting with their health care providers. Regardless of the efficacy of such natural products for alleviating menopausal discomfort, hops, licorice species, and their bioactive constituents such as 8-PN, XH, LigC, LigF, 8-PA, and LicA have been shown to activate various chemoprevention targets (8,10). It was shown that hops and its bioactive compound 6-PN enhanced CYP1A1 enzyme activity that leads to benign estrogen metabolism and is considered a detoxifying chemoprevention pathway (15). In addition, hops and its abundant electrophilic compound, XH, were shown to activate NRF2 dependent antioxidant responses including the detoxification enzymes glutathione-S-transferase (GST) and NADPH:quinone oxidoreductase 1 (NQO1) *in vitro* and *in vivo* (11). Additionally, hops and its bioactive compounds were reported to have aromatase inhibitory effects *in vitro* (45–48). The licorice species, GI, and the unique bioactive chalcone of GI, LicA, were shown to suppress the levels of estrogen carcinogenic metabolites through the suppression of CYP1B1 *in vitro* and *in vivo* (12,16). The three licorice species GG, GU, and GI as well as their compounds LigC and LicA induced NQO1 activity *in vitro*, while GG induced NQO1 activity in the mammary tissue and GI and LicA increased NQO1 activity in the liver *in vivo* (13,16). LigC from licorice was also reported to have aromatase inhibitory potential, *in vitro* (49,50). In addition, GG, GI, and their bioactive compounds, LigC and LicA, have shown anti-inflammatory potential through the suppression of iNOS activity in murine macrophages (12).

Collectively, these findings suggest there might be a multitude of chemopreventive pathways modulated by hops, licorice species, and their bioactive compounds, making them suitable for further studies. Aromatase inhibition is a known chemopreventive strategy for high-risk postmenopausal women. Establishing such an effect for hops, licorice, and their bioactive compounds would lead to an alternative breast cancer prevention strategy with desired efficacy and potentially greater acceptability compared to the available interventions.

Estrogen (E_1_/E_2_) production in the breast takes place in the pre-adipocyte fibroblasts of the stromal compartment of the breast (3). The rate of E_1_/E_2_ biosynthesis is mainly regulated through the activity of aromatase, encoded by the *CYP19A1* gene in humans. While protein expression is homologous across different tissues, its expression levels are controlled by the interaction of distinct transcription factors:, at least 10 different tissue specific and alternative promoters upstream of the *CYP19A1* gene are known, resulting in various expression levels of the aromatase enzyme (51). Alternate activation of promoter II (PII) and promoter I.3 (PI.3) instead of promoter I.4 (PI.4) can lead to excessive aromatase expression in the cancerous breast (51). In the cancer-free breast; however, activation of PI.4 by factors such as pro-inflammatory cytokines, which can be high in the postmenopausal breast, leads to elevated aromatase expression, resulting in excessive conversion of androgens to E_1_/E_2_ (41,52). Locally produced E_1_/E_2_ enter the epithelial compartment in a paracrine manner, fueling the proliferation of epithelial cells and the genotoxic estrogen metabolism leading to carcinogenesis (52). Compounds suppressing *CYP19A1* expression or directly inhibiting aromatase activity could potentially reduce local estrogen biosynthesis in the breast and serve as potential chemopreventive agents against breast cancer.

The present observations suggest that hops, GG, GU, and GI contain aromatase inhibiting constituents and exhibit varying levels of inhibitory potential (Figure 3). The highest inhibition potency observed with GI (Figure 3) compared to the other extracts suggests the presence of compound(s) with significant inhibitory potential in this extract. Data obtained with the pure compounds showed that the phytoestrogen such as LigF, which is common in all licorice species, and 8-PA, which is more abundant in GI, had aromatase inhibitory potencies at nanomolar concentrations (Table 1). The GI specific bioactives, LigC and LicA, exhibited only moderate inhibitory effects. Therefore, the observed inhibitory potency with GI could be associated with the presence of these constituents, LigF, 8-PA, LigC, and LicA, in addition to effects of other unknown constituents in the extract.

Compared to GI, hops exhibited a moderate aromatase inhibition (Figure 3), despite its potent phytoestrogen, 8-PN, exhibiting the highest aromatase inhibition effect compared to all the tested compounds (Figure 4A). This observation can be explained by the relatively low abundance of 8-PN in hops extract (0.33%), whereas its biogenetic precursor, XH, is a moderate aromatase inhibitor yet much more abundant constituent (32% of the extract). Conversion of XH to 8-PN can be excluded under the conditions of *in vitro* aromatase inhibition assays. However, when extract is ingested in vivo, more 8-PN will form via IX as intermediate which is in equilibrium with XH (Figure 2B), through the metabolic activities of cytochrome P450’s and gut microbiota, leading to potentially more pronounced aromatase inhibitory effects with hops extract *in vivo*.

Previous studies have shown estrogenic effects for 8-PN from hops, LigF, and 8-PA from licorice (Table 1) (30,37). The current study showed that these phytoestrogens were better aromatase inhibitors compared to the other tested bioactive compounds from hops and licorice species (Table 1). The potent phytoestrogen, 8-PN, also proved to be the most potent aromatase inhibitor (Table 1). Therefore, binding of these phytoestrogens to the aromatase binding pocket was compared to known representatives of two classes of aromatase inhibitors, letrozole (non-steroidal) and exemestane (steroidal). These were evaluated computationally, using in silico models of aromatase (Figure 5) (26,30,37). These calculations suggest that the potent aromatase inhibitors, 8-PN from hops, and LigF and 8-PA from licorice, bind in the aromatase binding pocket similarly to the known inhibitors, letrozole and exemestane.

While both exemestane and letrozole block the binding of androstenedione via their binding to residues within aromatase’s binding pocket, exemestane irreversibly binds, while letrozole reversibly coordinates with heme iron and residues (39). The compounds with high binding affinity bind in a similar location to letrozole and exemestane (Figure 5). In their most energetically favorable conformation, 8-PN and LigF bind to the heme complex (Figure 5). Past studies found that removal or blocking of the C4 carbonyl present in these compounds greatly reduced binding affinity, further suggesting the importance of heme interaction (53–55).

Although letrozole’s heme interaction is known, the conformation of the rest of the molecule is less well-established. In this study, in its most favorable position, one of letrozole’s nitrile adjacent rings is located in a hydrophobic channel of the binding pocket (Figure S1B) as documented in literature (56). The hydrophobic prenyl group of the best binder, 8-PN, and the third best binder, 8-PA, extends in the area of this same hydrophobic channel (Figure 5, Figure S3). Likewise, past aromatase inhibition studies have found C8 prenylation of naringenin (8-PN) decreased the IC_50_ more than10-fold, while C6 prenylation led to poor aromatase binding affinity (10,57). When combined with the results of the current study, these findings suggest that 8-PN and 8-PA may bind well due to their prenyl group falling within the hydrophobic channel when heme-binding occurs. Moreover, in its most energetically favorable conformation, 8-PN directly binds to heme while 8-PA adopts a less energetically favorable position to achieve heme binding (data not shown), a potential explanation for the significant difference in binding affinities of these two structurally similar compounds.

The compounds with microM IC_50_ values, XH, 6-PN, LigC, and LicA, neither bound within the binding pocket, nor interacted directly with heme or had groups in energetically unfavorable positions. XH (data not shown) did not fall fully within the binding pocket when bound in its most energetically favorable position. Although falling within the binding pocket, LicA (Figure S2A) and LigC (data not shown) did not directly bind to the heme group, whereas 6-PN did. However, the hydrophobic prenyl group of 6-PN was located an unfavorable position within a hydrophilic region (Figure S3A). While these computational results should be confirmed in a future structure-activity relationship study, they suggest that the phytoestrogens 8-PN from hops and LigF and 8-PA from licorice species have the potential to act as natural aromatase inhibitors.

In addition to the direct inhibition of the enzyme activity, limiting estrogen production could also be achieved by suppressing the expression of the aromatase gene, *CYP19A1*, which will in turn result in lower active enzyme levels and limited E_1_/E_2_ production. Aromatase expression in the cancer-free breast is regulated through the promoter PI.4 which can be upregulated by pro-inflammatory cytokines generated by the resident macrophages in the breast microenvironment (58). Studies have shown that menopause is associated with enhanced adiposity and increased inflammation in the breast which leads to increased aromatase expression and estrogen exposure (3). Data from this study showed that the baseline levels of *CYP19A1* in control microstructures (obtained from high-risk postmenopausal breast tissue) were high. Therefore, the effects of hops and its bioactive compounds 8-PN, 6-PN, XH, as well as licorice species, and their bioactive compounds LigF, LigC, LicA, and 8-PA on the expression of *CYP19A1* were evaluated. Microstructures contain the variety of cell types in the breast, maintain the protein expression patterns, and the architectural features of the original tissue (34). Therefore, the effects of the treatments on the breast can be defined in a more realistic manner. In this study, microstructures were obtained from the fresh surgically removed breast tissue of high-risk postmenopausal women undergoing bilateral mastectomy due to incident unilateral breast cancer. Among the tested extracts and bioactive compounds were hops and GI as well as their bioactive compounds. The results showed that exposure to hops, 8-PN, GI, LigF, and LicA for 24 h did not affect the morphology of the microstructures in culture; however, 8-PN from hops, GI extract, LigF, and LicA decreased the expression of the *CYP19A1* significantly (Figure 6). These findings support the conclusion that these natural products might have effects on regulating the promoter PI.4 of aromatase. As this promoter is particularly important in cancer-free breast tissue, its regulation in the unaffected breast microstructures could be directly relevant to cancer prevention. Further studies are needed to define how the studies natural products modulate regulatory factors including cytokines responsible for regulating PI.4.

**Figure 6:**
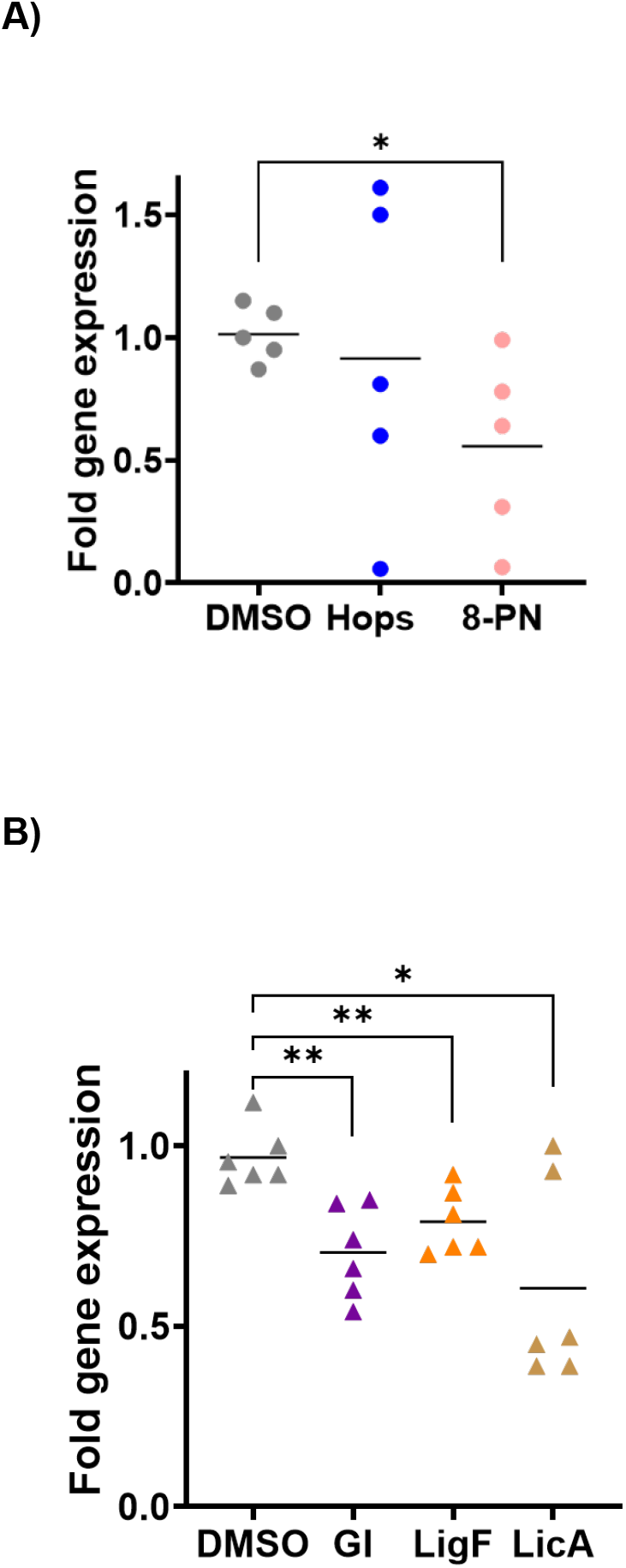
Hops and GI suppress the expression of CYP19A1 mRNA in high-risk breast tissue microstructures. Fresh surgically removed unaffected contralateral breast tissue from postmenopausal women undergoing double mastectomy due to unilateral breast cancer were processed to make microstructures using collagenase I, before culturing and treating them for 24 h with A) hops (5 μg/mL), 8-PN (10 μM) vs. B) GI (5 μg/mL), LigF (5 μM), LicA (5 μM). Subsequently, RNA extraction and qPCR of CYP19A1 was performed. The expression of CYP19A1 mRNA was calculated relative to the housekeeping gene RPLP0 and normalized to DMSO control. Data represents mean ± SD of three independent observations (p < 0.05).

Overall, the data suggest that widely used women’s health botanicals such as hops, GG, GU, GI (Figure 3), and their phytoestrogens 8-PN, LigF, and 8-PA (Figure 2, Figure 4) do have potential aromatase inhibition effects. 8-PN, GI, LigF, and LicA also suppress aromatase expression (Figure 6) in the high-risk postmenopausal women’s breast microstructures. Thus, hops, licorice, and their bioactive compounds might have the potential to limit estrogen exposure in the postmenopausal breast, which is an important goal in breast cancer prevention. Future studies using high-risk breast tissue of additional subjects and *in vivo* evaluations are under way to further validate these observations.

## Conclusions

Considering the study outcomes in conjunction with previously reported chemopreventive properties such as the suppression of estrogen genotoxic metabolism, induction of detoxification pathways, and anti-inflammatory effects with these natural products, it can be concluded that the studied botanical interventions are promising and worth further investigations towards defining as alternative risk reducing interventions for postmenopausal breast cancer (8,10–16). The results also provide the basis for botanical standardization efforts aimed at enriching extracts for cancer prevention purpose, through the innovative DESIGNER methodology, which refers to designing extracts that differ in one (or very) few compounds or a group of congeners (compounds structurally and pharmacologically relevant), while otherwise having (near-)identical complex chemical composition (59–61). Considering that natural products do not share the barrier of intervention acceptability that is the major obstacle with presently available prevention drugs, the presented evidence of aromatase-based mechanisms of action supports the risk reducing potential of these natural interventions (2). The outcomes of continued investigations can provide postmenopausal women and their health care providers with evidence-based alternative intervention options. The presented data also assists with making educated decisions or giving advice about botanical dietary supplements for postmenopausal women.

## Acknowledgements

We thank Dr. Liang Zhao, LICP, CAS, for a generous gift of *G. inflata* plant material. We also thank Natalie Pulliam for coordinating the clinical specimen collection and consenting the subjects. We are thankful to the surgical pathology laboratory at Northwestern Memorial Hospital and Kristy Skurauskis for characterizing and providing the surgical tissue used in this study.

## Abbreviations

AKR1C3: Aldoketoreductase 1C3
AI: Aromatase inhibitor
CYP: Cytochrome P450
E_2_: 17β-Estradiol
ER+: Estrogen receptor positive
E_1_: Estrone
Ex/Em: Excitation/Emission
GG: *Glycyrrhiza glabra*
GI: *Glycyrrhiza inflata*
GU: *Glycyrrhiza uralensis*
G6PDH: Glucose-6-phosphate dehydrogenase
GST: Glutathione-S-transferase
HBSS: Hank’s balanced salt solution
HFC: 7-Hydroxy-4-trifluoromethyl coumarin
IRB: Institutional Review Board
IDT: Integrated DNA Technologies
LigC: Isoliquiritigenin
IX: Isoxanthohumol
LicA: Licochalcone A
LigF: Liquiritigenin
MFC: 7-methoxy-4-trifluoromethyl coumarin
NADPH: Nicotineamide adenine dinucleotide phosphate
NQO1: NADPH:quinone oxidoreductase 1
6-PN: 6-prenylnaringenin
8-PA: 8-prenylapigenin
8-PN: 8-prenylnaringenin
PII: Promoter II
PI.3: promoter I.3
PI.4: Promoter I.4
SERM: Selective estrogen receptor modulators
XH: Xanthohumol

## Bibliography

1. Bray F, Ferlay J, Soerjomataram I, Siegel RL, Torre LA, Jemal A. Global cancer statistics 2018: GLOBOCAN estimates of incidence and mortality worldwide for 36 cancers in 185 countries. CA Cancer J Clin 2018;68:394–424 doi 10.3322/caac.21492.

2. Sun YS, Zhao Z, Yang ZN, Xu F, Lu HJ, Zhu ZY, et al. Risk Factors and Preventions of Breast Cancer. Int J Biol Sci 2017;13:1387–97 doi 10.7150/ijbs.21635.

3. Brown KA, Iyengar NM, Zhou XK, Gucalp A, Subbaramaiah K, Wang H, et al. Menopause Is a Determinant of Breast Aromatase Expression and Its Associations With BMI, Inflammation, and Systemic Markers. J Clin Endocrinol Metab 2017;102:1692–701 doi 10.1210/jc.2016-3606.

4. Nelson LR, Bulun SE. Estrogen production and action. J Am Acad Dermatol 2001;45:S116–24 doi 10.1067/mjd.2001.117432.

5. Trivedi MS, Coe AM, Vanegas A, Kukafka R, Crew KD. Chemoprevention Uptake among Women with Atypical Hyperplasia and Lobular and Ductal Carcinoma In Situ. Cancer Prev Res (Phila) 2017;10:434–41 doi 10.1158/1940-6207.CAPR-17-0100.

6. Avvaru SP, Noolvi MN, Aminbhavi TM, Chkraborty S, Dash A, Shukla SS. Aromatase Inhibitors Evolution as Potential Class of Drugs in the Treatment of Postmenopausal Breast Cancer Women. Mini Rev Med Chem 2018;18:609–21 doi 10.2174/1389557517666171101100902.

7. Hum S, Wu M, Pruthi S, Heisey R. Physician and Patient Barriers to Breast Cancer Preventive Therapy. Curr Breast Cancer Rep 2016;8:158–64 doi 10.1007/s12609-016-0216-5.

8. Dietz BM, Hajirahimkhan A, Dunlap TL, Bolton JL. Botanicals and Their Bioactive Phytochemicals for Women’s Health. Pharmacol Rev 2016;68:1026–73 doi 10.1124/pr.115.010843.

9. Hajirahimkhan A, Dietz BM, Bolton JL. Botanical modulation of menopausal symptoms: mechanisms of action? Planta Med 2013;79:538–53 doi 10.1055/s-0032-1328187.

10. Bolton JL, Dunlap TL, Hajirahimkhan A, Mbachu O, Chen SN, Chadwick L, et al. The Multiple Biological Targets of Hops and Bioactive Compounds. Chem Res Toxicol 2019;32:222–33 doi 10.1021/acs.chemrestox.8b00345.

11. Dietz BM, Hagos GK, Eskra JN, Wijewickrama GT, Anderson JR, Nikolic D, et al. Differential regulation of detoxification enzymes in hepatic and mammary tissue by hops *(Humulus lupulus)* in vitro and in vivo. Mol Nutr Food Res 2013;57:1055–66 doi 10.1002/mnfr.201200534.

12. Dunlap TL, Wang S, Simmler C, Chen SN, Pauli GF, Dietz BM, et al. Differential Effects of *Glycyrrhiza* Species on Genotoxic Estrogen Metabolism: Licochalcone A Downregulates P450 1B1, whereas Isoliquiritigenin Stimulates It. Chem Res Toxicol 2015;28:1584–94 doi 10.1021/acs.chemrestox.5b00157.

13. Hajirahimkhan A, Simmler C, Dong H, Lantvit DD, Li G, Chen SN, et al. Induction of NAD(P)H:Quinone Oxidoreductase 1 (NQO1) by *Glycyrrhiza* Species Used for Women’s Health: Differential Effects of the Michael Acceptors Isoliquiritigenin and Licochalcone A. Chem Res Toxicol 2015;28:2130–41 doi 10.1021/acs.chemrestox.5b00310.

14. Hemachandra LP, Madhubhani P, Chandrasena R, Esala P, Chen SN, Main M, et al. Hops (Humulus lupulus) inhibits oxidative estrogen metabolism and estrogen-induced malignant transformation in human mammary epithelial cells (MCF-10A). Cancer Prev Res (Phila) 2012;5:73–81 doi 10.1158/1940-6207.CAPR-11-0348.

15. Wang S, Dunlap TL, Howell CE, Mbachu OC, Rue EA, Phansalkar R, et al. Hop (Humulus lupulus L.) Extract and 6-Prenylnaringenin Induce P450 1A1 Catalyzed Estrogen 2-Hydroxylation. Chem Res Toxicol 2016;29:1142–50 doi 10.1021/acs.chemrestox.6b00112.

16. Wang S, Dunlap TL, Huang L, Liu Y, Simmler C, Lantvit DD, et al. Evidence for Chemopreventive and Resilience Activity of Licorice: *Glycyrrhiza Glabra* and *G*. *Inflata* Extracts Modulate Estrogen Metabolism in ACI Rats. Cancer Prev Res (Phila) 2018;11:819–30 doi 10.1158/1940-6207.CAPR-18-0178.

17. Cho YC, Kim HJ, Kim YJ, Lee KY, Choi HJ, Lee IS, et al. Differential anti-inflammatory pathway by xanthohumol in IFN-gamma and LPS-activated macrophages. Int Immunopharmacol 2008;8:567–73 doi 10.1016/j.intimp.2007.12.017.

18. Ferk F, Huber WW, Filipic M, Bichler J, Haslinger E, Misik M, et al. Xanthohumol, a prenylated flavonoid contained in beer, prevents the induction of preneoplastic lesions and DNA damage in liver and colon induced by the heterocyclic aromatic amine amino-3-methyl-imidazo[4,5-f]quinoline (IQ). Mutat Res 2010;691:17–22 doi 10.1016/j.mrfmmm.2010.06.006.

19. Henderson MC, Miranda CL, Stevens JF, Deinzer ML, Buhler DR. In vitro inhibition of human P450 enzymes by prenylated flavonoids from hops, Humulus lupulus. Xenobiotica 2000;30:235–51 doi 10.1080/004982500237631.

20. Jiang CH, Sun TL, Xiang DX, Wei SS, Li WQ. Anticancer Activity and Mechanism of Xanthohumol: A Prenylated Flavonoid From Hops (Humulus lupulus L.). Front Pharmacol 2018;9:530 doi 10.3389/fphar.2018.00530.

21. Monteiro R, Calhau C, Silva AO, Pinheiro-Silva S, Guerreiro S, Gartner F, et al. Xanthohumol inhibits inflammatory factor production and angiogenesis in breast cancer xenografts. J Cell Biochem 2008;104:1699–707 doi 10.1002/jcb.21738.

22. Pichler C, Ferk F, Al-Serori H, Huber W, Jager W, Waldherr M, et al. Xanthohumol Prevents DNA Damage by Dietary Carcinogens: Results of a Human Intervention Trial. Cancer Prev Res (Phila) 2017;10:153–60 doi 10.1158/1940-6207.CAPR-15-0378.

23. Cuendet M, Guo J, Luo Y, Chen S, Oteham CP, Moon RC, et al. Cancer chemopreventive activity and metabolism of isoliquiritigenin, a compound found in licorice. Cancer Prev Res (Phila) 2010;3:221–32 doi 10.1158/1940-6207.CAPR-09-0049.

24. Bode AM, Dong Z. Chemopreventive Effects of Licorice and Its Components. Curr Pharmacol Rep 2015;1:60–71 doi 10.1007/s40495-014-0015-5.

25. Chadwick LR, Nikolic D, Burdette JE, Overk CR, Bolton JL, van Breemen RB, et al. Estrogens and congeners from spent hops (Humulus lupulus). J Nat Prod 2004;67:2024–32 doi 10.1021/np049783i.

26. Krause E, Yuan Y, Hajirahimkhan A, Dong H, Dietz BM, Nikolic D, et al. Biological and chemical standardization of a hop (Humulus lupulus) botanical dietary supplement. Biomed Chromatogr 2014;28:729–34 doi 10.1002/bmc.3177.

27. Dietz BM, Chen SN, Alvarenga RFR, Dong H, Nikolic D, Biendl M, et al. DESIGNER Extracts as Tools to Balance Estrogenic and Chemopreventive Activities of Botanicals for Women’s Health. J Nat Prod 2017;80:2284–94 doi 10.1021/acs.jnatprod.7b00284.

28. van Breemen RB, Yuan Y, Banuvar S, Shulman LP, Qiu X, Alvarenga RF, et al. Pharmacokinetics of prenylated hop phenols in women following oral administration of a standardized extract of hops. Mol Nutr Food Res 2014;58:1962–9 doi 10.1002/mnfr.201400245.

29. Simmler C, Anderson JR, Gauthier L, Lankin DC, McAlpine JB, Chen SN, et al. Metabolite Profiling and Classification of DNA-Authenticated Licorice Botanicals. J Nat Prod 2015;78:2007–22 doi 10.1021/acs.jnatprod.5b00342.

30. Hajirahimkhan A, Simmler C, Yuan Y, Anderson JR, Chen SN, Nikolic D, et al. Evaluation of estrogenic activity of licorice species in comparison with hops used in botanicals for menopausal symptoms. PLoS One 2013;8:e67947 doi 10.1371/journal.pone.0067947.

31. Kondo K, Shiba M, Yamaji H, Morota T, Zhengmin C, Huixia P, et al. Species identification of licorice using nrDNA and cpDNA genetic markers. Biol Pharm Bull 2007;30:1497–502 doi 10.1248/bpb.30.1497.

32. Pauli GF, Chen SN, Simmler C, Lankin DC, Godecke T, Jaki BU, et al. Importance of purity evaluation and the potential of quantitative (1)H NMR as a purity assay. J Med Chem 2014;57:9220–31 doi 10.1021/jm500734a.

33. Tanos T, Sflomos G, Echeverria PC, Ayyanan A, Gutierrez M, Delaloye JF, et al. Progesterone/RANKL is a major regulatory axis in the human breast. Sci Transl Med 2013;5:182ra55 doi 10.1126/scitranslmed.3005654.

34. Cartaxo AL, Estrada MF, Domenici G, Roque R, Silva F, Gualda EJ, et al. A novel culture method that sustains ERalpha signaling in human breast cancer tissue microstructures. J Exp Clin Cancer Res 2020;39:161 doi 10.1186/s13046-020-01653-4.

35. Bhatnagar AS, Hausler A, Schieweck K, Lang M, Bowman R. Highly selective inhibition of estrogen biosynthesis by CGS 20267, a new non-steroidal aromatase inhibitor. J Steroid Biochem Mol Biol 1990;37:1021–7 doi 10.1016/0960-0760(90)90460-3.

36. Kondo K, Shiba M, Nakamura R, Morota T, Shoyama Y. Constituent properties of licorices derived from Glycyrrhiza uralensis, G. glabra, or G. inflata identified by genetic information. Biol Pharm Bull 2007;30:1271–7 doi 10.1248/bpb.30.1271.

37. Hajirahimkhan A, Mbachu O, Simmler C, Ellis SG, Dong H, Nikolic D, et al. Estrogen Receptor (ER) Subtype Selectivity Identifies 8-Prenylapigenin as an ERbeta Agonist from *Glycyrrhiza inflata* and Highlights the Importance of Chemical and Biological Authentication. J Nat Prod 2018;81:966–75 doi 10.1021/acs.jnatprod.7b01070.

38. Overk CR, Guo J, Chadwick LR, Lantvit DD, Minassi A, Appendino G, et al. In vivo estrogenic comparisons of Trifolium pratense (red clover) Humulus lupulus (hops), and the pure compounds isoxanthohumol and 8-prenylnaringenin. Chem Biol Interact 2008;176:30–9 doi 10.1016/j.cbi.2008.06.005.

39. Hong Y, Yu B, Sherman M, Yuan YC, Zhou D, Chen S. Molecular basis for the aromatization reaction and exemestane-mediated irreversible inhibition of human aromatase. Mol Endocrinol 2007;21:401–14 doi 10.1210/me.2006-0281.

40. Dunneram Y, Greenwood DC, Cade JE. Diet, menopause and the risk of ovarian, endometrial and breast cancer. Proc Nutr Soc 2019;78:438–48 doi 10.1017/S0029665118002884.

41. Iyengar NM, Morris PG, Zhou XK, Gucalp A, Giri D, Harbus MD, et al. Menopause is a determinant of breast adipose inflammation. Cancer Prev Res (Phila) 2015;8:349–58 doi 10.1158/1940-6207.CAPR-14-0243.

42. Maresso KC, Tsai KY, Brown PH, Szabo E, Lippman S, Hawk ET. Molecular cancer prevention: Current status and future directions. CA Cancer J Clin 2015;65:345–83 doi 10.3322/caac.21287.

43. Martinez KA, Fagerlin A, Witteman HO, Holmberg C, Hawley ST. What Matters to Women When Making Decisions About Breast Cancer Chemoprevention? Patient 2016;9:149–59 doi 10.1007/s40271-015-0134-z.

44. Colditz GA, Bohlke K. Priorities for the primary prevention of breast cancer. CA Cancer J Clin 2014;64:186–94 doi 10.3322/caac.21225.

45. Monteiro R, Becker H, Azevedo I, Calhau C. Effect of hop (Humulus lupulus L.) flavonoids on aromatase (estrogen synthase) activity. J Agric Food Chem 2006;54:2938–43 doi 10.1021/jf053162t.

46. Monteiro R, Faria A, Azevedo I, Calhau C. Modulation of breast cancer cell survival by aromatase inhibiting hop (Humulus lupulus L.) flavonoids. J Steroid Biochem Mol Biol 2007;105:124–30 doi 10.1016/j.jsbmb.2006.11.026.

47. van Meeuwen JA, Nijmeijer S, Mutarapat T, Ruchirawat S, de Jong PC, Piersma AH, et al. Aromatase inhibition by synthetic lactones and flavonoids in human placental microsomes and breast fibroblasts--a comparative study. Toxicol Appl Pharmacol 2008;228:269–76 doi 10.1016/j.taap.2007.12.007.

48. Solak KA, Wijnolts FMJ, Nijmeijer SM, Blaauboer BJ, van den Berg M, van Duursen MBM. Excessive levels of diverse phytoestrogens can modulate steroidogenesis and cell migration of KGN human granulosa-derived tumor cells. Toxicol Rep 2014;1:360–72 doi 10.1016/j.toxrep.2014.06.006.

49. Ye L, Gho WM, Chan FL, Chen S, Leung LK. Dietary administration of the licorice flavonoid isoliquiritigenin deters the growth of MCF-7 cells overexpressing aromatase. Int J Cancer 2009;124:1028–36 doi 10.1002/ijc.24046.

50. Luo L, Shen L, Sun F, Ma Z. Immunoprecipitation coupled with HPLC-MS/MS to discover the aromatase ligands from Glycyrrhiza uralensis. Food Chem 2013;138:315–20 doi 10.1016/j.foodchem.2012.10.043.

51. Zhao H, Zhou L, Shangguan AJ, Bulun SE. Aromatase expression and regulation in breast and endometrial cancer. J Mol Endocrinol 2016;57:R19–33 doi 10.1530/JME-15-0310.

52. Wang X, Simpson ER, Brown KA. Aromatase overexpression in dysfunctional adipose tissue links obesity to postmenopausal breast cancer. J Steroid Biochem Mol Biol 2015;153:35–44 doi 10.1016/j.jsbmb.2015.07.008.

53. Kao YC, Zhou C, Sherman M, Laughton CA, Chen S. Molecular basis of the inhibition of human aromatase (estrogen synthetase) by flavone and isoflavone phytoestrogens: A site-directed mutagenesis study. Environ Health Perspect 1998;106:85–92 doi 10.1289/ehp.9810685.

54. Mojaddami A, Sakhteman A, Fereidoonnezhad M, Faghih Z, Najdian A, Khabnadideh S, et al. Binding mode of triazole derivatives as aromatase inhibitors based on docking, protein ligand interaction fingerprinting, and molecular dynamics simulation studies. Res Pharm Sci 2017;12:21–30 doi 10.4103/1735-5362.199043.

55. Neves MA, Dinis TC, Colombo G, Sa e Melo ML. Combining computational and biochemical studies for a rationale on the anti-aromatase activity of natural polyphenols. ChemMedChem 2007;2:1750–62 doi 10.1002/cmdc.200700149.

56. Loge C, Le Borgne M, Marchand P, Robert JM, Le Baut G, Palzer M, et al. Three-dimensional model of cytochrome P450 human aromatase. J Enzyme Inhib Med Chem 2005;20:581–5 doi 10.1080/14756360500220574.

57. Nielsen AJ, McNulty J. Polyphenolic natural products and natural product-inspired steroidal mimics as aromatase inhibitors. Med Res Rev 2019;39:1274–93 doi 10.1002/med.21536.

58. Gerard C, Brown KA. Obesity and breast cancer-Role of estrogens and the molecular underpinnings of aromatase regulation in breast adipose tissue. Mol Cell Endocrinol 2018;466:15–30 doi 10.1016/j.mce.2017.09.014.

59. Yu Y, Simmler C, Kuhn P, Poulev A, Raskin I, Ribnicky D, et al. The DESIGNER Approach Helps Decipher the Hypoglycemic Bioactive Principles of Artemisia dracunculus (Russian Tarragon). J Nat Prod 2019;82:3321–9 doi 10.1021/acs.jnatprod.9b00548.

60. Malca Garcia GR, Friesen JB, Liu Y, Nikolic D, Lankin DC, McAlpine JB, et al. Preparation of DESIGNER extracts of red clover (Trifolium pratense L.) by centrifugal partition chromatography. J Chromatogr A 2019;1605:360277 doi 10.1016/j.chroma.2019.05.057.

61. Friesen JB, Liu Y, Chen SN, McAlpine JB, Pauli GF. Selective Depletion and Enrichment of Constituents in “Curcumin” and Other Curcuma longa Preparations. J Nat Prod 2019;82:621–30 doi 10.1021/acs.jnatprod.9b00020.

