## Supplementary data for "Inhibition of Aromatase by Hops, Licorice Species, and their bioactive compounds in Postmenopausal Breast Tissue"

† In memory of Dr. Judy L. Bolton.

**Correspondence**

Dr. Atieh Hajirahimkhan, Ph.D., American Cancer Society postdoctoral fellow, Robert. H. Lurie Comprehensive Cancer Center, Department of Surgery, Feinberg School of Medicine, Northwestern University, 303 E. Superior, 4-220, Chicago, IL. 60611., Phone: 312-503-2109

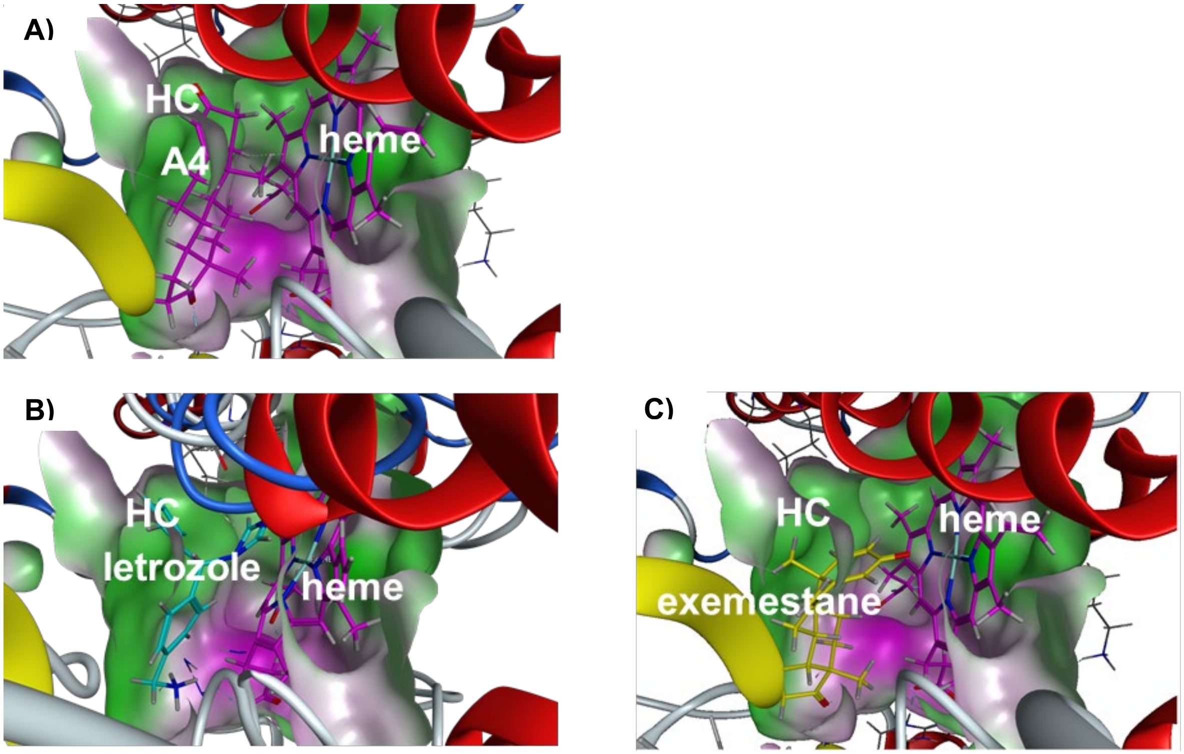


**Figure S1:** A) Androstendione (A4), B) letrozole, ~~or~~ and C) exemestane in the binding pocket of aromatase with heme present. Green surfaces represent~~s~~ hydrophobic regions; purple surfaces represent~~s~~ hydrophilic regions of the binding pocket. HC is the hydrophobic channel as discussed in the main text~~paper~~.

A)

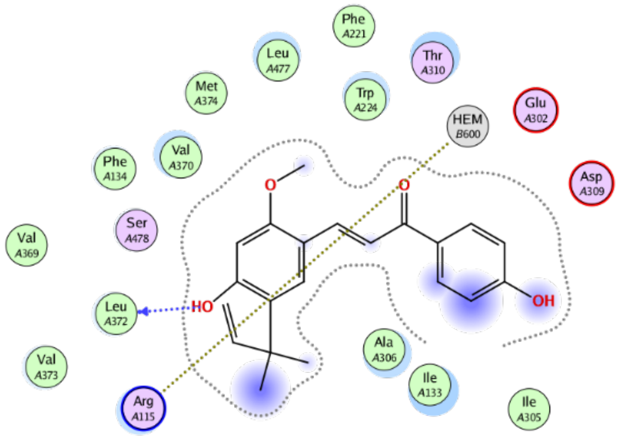


B)

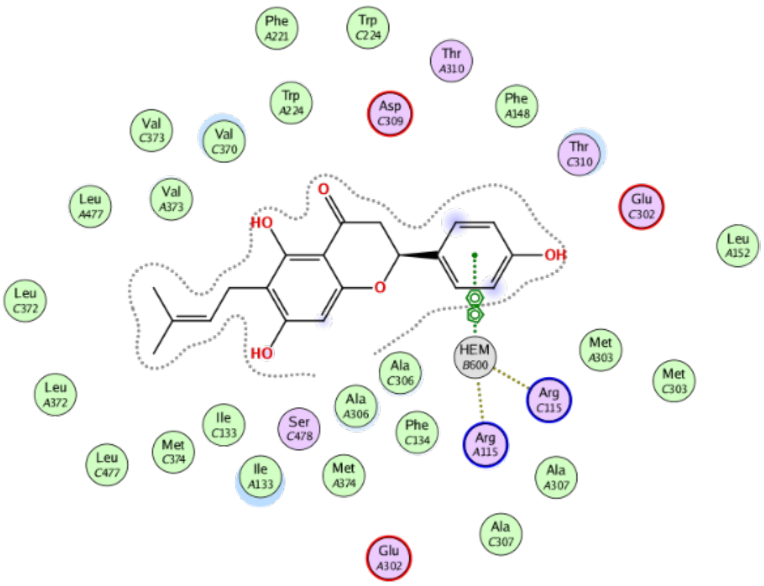


**Figure S2:** Whereas LicA does not directly bind to heme (HEM) in the binding site of aromatase, 6-PN does: A) 2D interaction diagram of the most energetically favorable conformation of LicA; B) 2D overlay interaction diagram of top two most energetically favorable binding conformations of 6-PN bound to aromatase (3EQM).

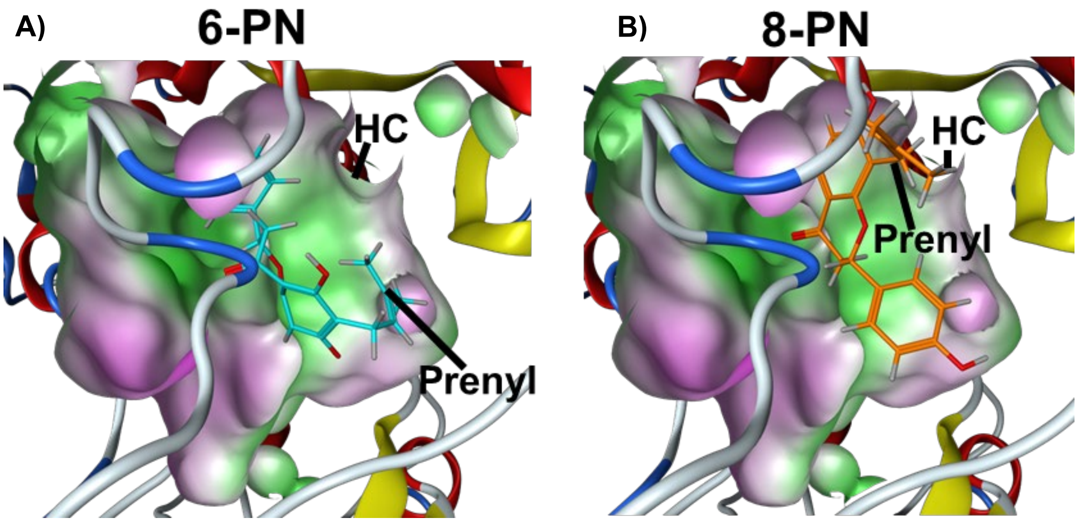


**Figure S3:** While the prenyl group o 6-PN does not fall within the hydrophobic channel (HC) of the aromatase protein, the same moiety of 8-PN does: A) 6-PN (light blue) vs. B) 8-PN (orange) docked into aromatase (ribbon)-heme (removed from picture) complex in its most energetically favorable conformation with hydrophobicity surface. Green surfaces are the hydrophobic region of binding pocket; purple surfaces are the hydrophilic regions.
